# Where does it go? The fate of thiocyanate in the aquarium water and blood plasma of *Amphiprion clarkii* after exposure to cyanide

**DOI:** 10.1101/2020.05.04.076430

**Authors:** J. Alexander Bonanno, Nancy E. Breen, Michael F. Tlusty, Lawrence J. Andrade, Andrew L. Rhyne

## Abstract

The illegal practice of cyanide fishing continues to damage coral reef ecosystems throughout the Indo-Pacific. To combat this destructive fishing method, a simple, reliable test to detect whether or not a fish has been captured using cyanide (CN) is needed. This study analyzed the toxicokinetics of acute, pulsed CN exposure as well as chronic exposure to thiocyanate (SCN), the major metabolite of CN, in the clownfish species, *Amphiprion clarkii*. Fish were pulse exposed to 50 ppm CN for 20 or 45 seconds or chronically exposed to 100 ppm SCN for 12 days. Blood plasma levels of SCN were measured following derivatization to SCN-bimane using an Acquity UPLC I-Class and Q-Exactive hybrid Quadrupole-Orbitrap HRAM mass spectrometer or directly by HPLC-UV. After exposure to CN, depending on the duration of exposure, SCN plasma levels reached a maximum concentration (300–470 ppb) 0.13–0.17 days after exposure, had a 0.1 to 1.2 day half-life, and often did not return to baseline levels. The half-life of plasma SCN after direct exposure to SCN was found to be 0.13 days, similar to the CN exposure, and that SCN in the holding water would often drop below detection. Finally, we observed that when a fish, never exposed to SCN, was placed in aquarium water spiked with SCN, there was a steady decrease in aqueous SCN concentration over 24 hours until it could no longer be detected. This pattern was repeated with a second sequential dose. These results demonstrate that *A. clarkii* do not excrete SCN after CN exposure, but in fact can absorb low concentrations of SCN from water, refuting several publications. It appears that *A. clarkii* exhibit a classic two compartment model where SCN is rapidly eliminated from the blood plasma and is distributed throughout the tissue but not excreted in their urine. This study demonstrates that SCN may be used as a marker of CN exposure only if fish are tested shortly after exposure. There is species specific variability in response to CN, and studies of other taxa need to be performed before this test can be deployed in the field.

## INTRODUCTION

Coral reef ecosystems are being stressed to their tipping point largely by climate change, but other anthropogenic activities are playing a role in their destruction including illegal fishing practices (Burke et al. 2011; Hughes et al. 2017). This is critical as coral reefs globally boast the greatest diversity of marine life anywhere (Reaka-Kudla 1997; Bruno & Selig 2007). The biodiversity found in coral reefs make them one of the most valuable ecosystems on our planet, yet they are also one of the most threatened (Hughes et al. 2017).

One of the risks posed to coral reefs is the illegal fishing practice of using cyanide (CN) to capture reef fish. This practice is most often used for either the marine aquarium trade (MAT) or the live reef food fish trade (LRFT) and primarily occurs in the Indo-Pacific region where 75 percent of all coral reefs are located (Rubec 1986; Barber & Pratt 1997; Graham 2001; Bruno & Selig 2007; Calado et al. 2014; Davis et al. 2017; Losada & Bersuder 2017). Cyanide fishing most often involves dissolving tablets of either potassium or sodium cyanide in a squirt bottle filled with seawater. The concentrated CN solution is then squirted onto the fish that inhabit the reef and in crevices in the reef where the fish hide. At sublethal doses, which likely vary depending on the size and species of the fish, CN temporally paralyzes the fish, immobilizing them for easy capture. Cyanide has been used as a stunning agent to collect fish for over 60 years (Lewis & Tarrant Jr 1960; Bellwood 1981; Pearce 2003; Frey 2013). Despite being illegal, this fishing practice continues to be used throughout the region. Fishermen rely on the middlemen, who provide cyanide, food, boats, and minimal pay in return for high volumes of cyanide-caught fish (Rubec et al. 2001). Policing against this fishing method is difficult, as there is no easy and definitive way to determine if a fish has been captured using CN. As a result, law enforcement is challenging if not impossible (reviewed by Losada & Bersuder, 2017).

The squirt bottle method of CN delivery makes it difficult for collectors to control the amount of solution dispensed, making dosing vary greatly. As CN is nonselective, and dosing is not precise, this method often kills both the targeted species and non-targeted species in the squirt zone (Frey 2013). Even when fish survive the exposure, there are reports of increased fish mortality post-capture associated with this fishing practice (Pyle 1993; Hall & Bellwood 1995; Cervino et al. 2003). Furthermore, stony corals, which are an important structural component of coral reef ecosystems, are often damaged from exposure to CN during the fishing process (Jones & Hoegh-Guldberg 1999; Cervino et al. 2003). Once the fish are stunned, fishermen also may break stony corals to gain access to the fish, increasing the long-term damage to reefs (Bruckner & Roberts 2008).

Attempts to curtail the use of CN as a capture method via post-capture testing have not been successful (Erdmann 1999; Dalabajan 2005). The Cyanide Detection Test (CDT), which uses a CN ion-selective electrode to measure CN found in tissues of fish, was developed by the International Marinelife Alliance (IMA) and the Philippines Bureau of Fisheries and Aquatic Resources (BFAR). It was adopted in the Philippines in the 1990s (Barber & Pratt 1997; Manipula et al. 2001). The CDT test was never validated for testing for CN collected fish, and it was suggested that the test was unreliable as an indicator of CN exposure primarily due to interferences resulting in false positives (Mak et al. 2005; Rubec et al. 2008; Balboa 2017). Bruckner and Roberts (2008) thoroughly documented the short comings of the IMA CDT. The test required lethal sampling, was time-consuming, labor-intensive, and was never properly field-tested and verified. Other CN detection methods have been proposed (Mak et al. 2005), but the rapid detoxification of CN is the major difficulty in detecting CN exposure. Any test for CN caught fish must be administered very soon, likely within hours, after exposure (Logue et al. 2010).

As an alternative to detecting CN directly, thiocyanate (SCN), the major metabolite of CN exposure, can be used as an indicator of CN exposure (Youso et al. 2012). In mammals, the CN detoxification pathway is well established. Two sulfur-transferases, rhodanese, and 3-mercaptopyruvate sulfurtransferase are responsible for catalyzing the formation of SCN from CN by sulfuration (Day et al. 2018), and SCN is eliminated in the urine by the kidneys. In marine fish, the method of elimination has not yet been determined, but Vaz et al. (2012) speculated a similar pathway to mammals. In a now refuted study, Vaz et al. (2012) claimed that a possible way to test for CN exposure in marine fish could be to detect SCN in the aquarium water of exposed fish. The authors claimed that the aquarium water contained the excreted metabolite SCN in sufficient quantity for detection. However, recovery of SCN from aquarium water as a test for CN exposure has never been replicated even though multiple labs have attempted to do so (Herz et al. 2016; Breen et al. 2018; Murray et al. 2020). In fact, a mass balance calculation demonstrated that the SCN levels reported by Vaz et al. (2012) were not possible because the CN dose needed to produce the reported amount of SCN is an order of magnitude higher than all the known LD50s in vertebrate species (Breen et al. 2018).

With the aquarium water test outside of the realm of possibility from a mass balance perspective, testing for SCN in bodily fluids of exposed marine fish is the next logical step. Indeed, elevated levels of SCN in the plasma of a marine fish after acute pulsed exposure to CN have been recently reported (Breen et al. 2019). SCN in the blood plasma of the laboratory cultured marine fish *Amphiprion ocellaris* was observed to be above control levels up to 41 days post exposure to CN. This study reported an initial fast elimination rate of SCN in blood plasma followed by a much slower elimination rate as evidenced by long residence times of low levels of SCN above that found in the controls. Breen et al. (2019) speculated that the observation of both a fast and slow elimination rate might be due to multiple elimination pathways in marine fish. These rates could also be governed by the availability of sulfur donors and the rate of diffusion from organs and tissues with limited blood flow (Day et al. 2018).

Breen et al. (2019) were the first to report the SCN concentration in the plasma of a marine fish following CN exposure. The report was for a single species and the species used was from an aquacultured stock with limited genetic diversity within the replicate fish. Over 2,300 documented reef fish taxa of varous sizes are traded in the MAT (Rhyne et al. 2017) and while the *A. ocellaris* work was a vital first step, the species dependent and fish size dependent variation of CN absorption and elimination is not known. Along with extending this work to other species, endogenous levels of SCN in marine fish must be known before any specific test that relies on SCN as a marker for CN exposure can be considered for widespread use.

In the present study, we report similar experiments to our previous CN toxicokinetic work using a congeneric clownfish species (*Amphiprion clarkii*), the same species used by Vaz et al. (2012). Here, *A. clarkii* were exposed to both CN and SCN and the rate of elimination of SCN from the blood plasma following exposure was determined by UPLC-MS and HPLC-UV. The toxicokinetics of SCN after pulsed exposure to CN is reported, as well as after chronic SCN exposure. In addition to testing the blood plasma, the aquarium water of *A. clarkii* was tested for SCN during depuration after chronic exposure to high concentrations (100 ppm) of SCN. With chronic, high-level direct exposure to the metabolite, the possibility of detection of SCN in aquarium water during depuration via bodily fluid excretion should be drastically enhanced.

## METHODS

### Test species and cyanide sources

Experiments were approved by the Roger Williams University Institute of Animal Use and Care Committee (Approval #R180820). *A. clarkii* of approximately 6-12 months of age were cultured in captivity at Roger Williams University or Sea & Reef Aquaculture, Franklin, ME, thereby ensuring they were not collected with or previously exposed to CN (Table 1). For all experiments, the water temperature was maintained by housing fish in a temperature-controlled room or placing buckets containing fish in a warm water bath both held at 25° C. All fish were fed pelletized food (Skretting Green Granule 1 mm) once per day unless otherwise noted. Water quality was maintained through daily water changes (100%) at a salinity of 30 ppt with light aeration.

**Table 1.** Average weight (± 1 S.D.) and sample size (n) of *Amphiprion clarkii* exposed to 50 ppm CN for either 20 or 45 seconds across 2 trials and 100 ppm for 12 day. There were two groups (small/large) of *Amphiprion clarkii* used for CN 2 trial however fish size did not affect SCN half-life.

| Trial | Exposure Time | Dose (ppm) | Time Sampled Post Exposure (days) | n | Average Weight (grams) |
| --- | --- | --- | --- | --- | --- |
| CN 1 | 20 s | 50 | 13 | 44 | 3.07 ± 0.62 |
| CN 1 | 45s | 50 | 13 | 48 | 3.54 ± 0.78 |
| CN 2 | 45 s | 50 | 72 | 41 | 12.84 ± 9.37<br>Small: 5.06 ± 1.33<br>Large: 21.0 ± 6.77 |
| SCN 1 | 12 d | 100 | 16 | 37 | 6.09 ± 2.72 |

### Exposures

For CN treatments, fish were exposed to a solution of 50 ppm CN (NaCN, Sigma 380970) for 20 s or 45 s across two separate trials (Table 1). Ten to 12 fish were placed in a basket, and immersed in the CN solution for the pre-determined time. Post-exposure, the fish were rinsed by transferring the basket containing the fish to two successive saltwater baths from the same source and at a salinity of 30 ppt. After rinsing, fish were housed by their exposure groups in round, 20 L polycarbonate tanks containing filtered saltwater with light aeration. Two trials of CN exposures were carried out. The first examined the SCN concentration in plasma for 13 days post exposure. Collection times were 1, 3, 6, 8, and 15 hours and 1, 2, 3, 7, 12 and 13 days post-exposure. The second served to determine how long the SCN concentration in the plasma of exposed fish remained above control levels. Collection times were 0.17, 0.5, 2, 7, 18, 50, and 72 days post-exposure. Plasma was also collected from a total of 22 control fish not exposed to CN across the two trials.

For the SCN treatment, fish were exposed to 100 ppm SCN (NaSCN, Sigma 467871) for 12 days (Table 1). Fish were housed in three separate round, 20 L polycarbonate tanks holding 12-13 fish each and 15 L of saltwater and fed daily. Complete water changes were performed daily. After 12 days, the fish were rinsed thoroughly by the treatment group in three consecutive saltwater baths in an attempt to remove all SCN from the surface of the fish. After rinsing, the fish were housed in the same groups as exposure in three separate 20 L polycarbonate tanks for depuration. Post-exposure tanks and fish were maintained as above. Once depuration was initiated, fish were sampled at 1, 2, 4, 8, and 16, hours and 1, 3 and 16 days. The aquarium water was also sampled over the first 48 hours as a pre-screen for the presence of excreted SCN.

### SCN in Aquarium Water

*Amphiprion clarkii* (n = 10) were exposed to 100 ppm SCN for 12 days in a 20 L polycarbonate tank as described above. To begin the depuration period, fish were rinsed individually in three consecutive saltwater baths. Rinse bath water was changed after each fish was rinsed. After rinsing, fish were placed in individual covered beakers containing 500 mL of saltwater. As a means to check for cross contamination, another 6 beakers were held in the same area, 3 containing non-exposed fish, and 3 without fish. Water samples (1 mL) were collected from each beaker before any fish were added, and then at 0, 2, 4, 8, 12, 24, 36, 48 and 72 hours after fish were added. Water changes were performed every 24 hours. Water samples were collected from each beaker before water changes during depuration. Fish were fed during exposure but not during depuration. After the final sampling at 72 hours, fish were bled and SCN blood plasma were measured as described below.

A second aquarium water sampling experiment was undertaken to further explore the results obtained in the first. Here, fish (n=10) with no known previous exposure to SCN, were placed in individual 500 mL beakers of saltwater spiked with SCN. In all cases, water samples were taken before spiking the water, after spiking the water with SCN and, 0, 2, 4, 8, 16, 24, (pre and post-SCN spike), and 48 hours after introducing fish to the beaker. Half the fish were bled at the 24 hour sampling before spiking the water again and the other half were bled after the final sampling at 48 hours. SCN blood plasma were measured as described below.

### Plasma extraction

At each time point fish were removed from the holding tanks for plasma collection (exact times are noted in supplemental material). Fish were heavily anesthetized with tricaine methanesulfonate (Western Chemical Inc., Ferndale, WA, USA) at a concentration of 200 ppm, buffered 2:1 with sodium bicarbonate in saltwater, and then dried and weighed on an analytical balance. Upon severance of the caudal peduncle with a #21 surgical blade, blood was collected in 40 mm heparinized microhematocrit tubes (Jorvet, Loveland, CO, USA) for fish weighing less than 4 g or 125 μL heparinized microcapillary blood collection tubes (RAM Scientific, Nashville, TN, USA) for fish greater than 4 g. Following blood collection fish were pithed. The 40 mm heparinized microhematocrit tubes were then centrifuged (ZipCombo Centrifuge, LW Scientific, Lawrenceville, GA, USA) at 3,000 rpm for two min followed by 6,000 rpm for five min to separate the red blood cells from the plasma and snapped at the plasma and red blood cell interface. The plasma was aspirated from the capillary tubes into pre-weighed 1.7 mL centrifuge tubes and then re-weighed to determine the mass of plasma collected. The 125 μL Micro-capillary blood collection tubes were spun at 12,000 rpm for 12 min. The top plasma layer was then pipetted into a pre-weighed 1.7 mL centrifuge tube which was then reweighed to determine mass. Plasma was stored in 1.7 mL centrifuge tubes at −80° C until they were analyzed.

### Plasma precipitation

Before plasma analysis, proteins were precipitated with cold HPLC grade acetonitrile (Sigma-Aldrich). Acetonitrile was added in the ratio of 1:5 (v/v) and the solution was vortexed for 20 s and then centrifuged at 12,000 rpm for 10 min. The supernatant was pipetted to a new 1.7 mL centrifuge tube and the acetonitrile was evaporated with warm nitrogen gas in an AutoVap (Zymark) at 70° C and then reconstituted in enough HPLC grade water (Sigma-Aldrich) for a 1:5 dilution and vortexed for 20 s.

### Thiocyanate analysis

Plasma SCN concentrations were analyzed following the method of Bhandari et al. (2014) and Breen et al. (2019). Diluted plasma (50 μL) was added to a 1.7 mL microcentrifuge tube, along with 25 μL of 4 mM MBB (Monobromobimane, Cayman Chemical 17097) in borate buffer (0.1 M, pH= 8) and 10 μL of the internal standard (200 ppb NaS^13^C^15^N) (Cambridge Isotope Labsyes). To minimize pipetting errors, larger volumes of the internal standard and the MBB were mixed in the appropriate volumes just prior to addition to the plasma and 35 μL of the resulting solution was added to the plasma. The plasma-MBB mixture was then heated to 70° C for 15 minutes to form the bimane-SCN complex. Standard solutions of SCN (10.0 ppb–25.0 ppm) were prepared in both HPLC grade water and in commercially available salmon plasma (MyBioSource inc., San Diego, CA, USA). Five-point calibration curves were prepared prior to all plasma analysis, such that the concentration of standards bracketed the expected plasma concentration.

Samples were analyzed for SCN-bimane concentration determination using a UPLC-MS system comprised of an Acquity UPLC^®^ I-Class (Waters Corp., Milford, MA) and Q-Exactive™ hybrid Quadrupole-Orbitrap™ high-resolution accurate mass (hram) mass spectrometer (Thermo Fisher Scientific, Waltham, MA). A Kinetex 1.7 μm XB-C18 100 Å 2.1 × 50 mm column held at 40° C was used for the gradient chromatographic separation (Phenomenex, Torrance, CA) with 10mm ammonium formate as mobile phase A and 10mm ammonium formate in methanol as mobile phase B. The flow rate was 0.25 ml/min, and gradient conditions were as follows: 10-100% B (0.00 – 3.00 min), 100% b (3.00 – 4.00 min), followed by 2. 00 min of re-equilibration time at initial conditions (total chromatographic run time 7.00 min). The Q-Exactive source conditions in esi negative mode were: sheath gas flow rate 55, auxiliary gas flow rate 15, sweep gas flow rate 2, spray voltage 4.50 kv, capillary temperature 300° C, s-lens rf level 55, auxiliary gas heater temperature 500°C. Tracefinder™ 3.2 was used for data acquisition and processing. Thermo Xcalibur™ 3.0 was also used for data processing.

The Q-Exactive acquisition method comprised a full-scan m/z 50-300 at 70,000 resolution. Analyte identity was established relative to a standard by scoring the following qualitative criteria using TraceFinder 3.2: retention time (rt ± 0.15 min), full-scan accurate mass (± 5 ppm window), full-scan isotope pattern (scores range 0-100, and a score of ≥ 70 was used as the positive cutoff). Analyte quantitation was performed using the peak area ratio from the full-scan extracted ion chromatograms (xics) of the analyte (248.04992) and its internal standard (250.05031); the xic mass window was the accurate mass ± 5 ppm in all cases.

Analyte validation and quantification was achieved by calculation of peak area ratio (analyte/internal standard) and subsequent concentration from a linear weighted (1/x) regression of calibration standards. To accept a calibration standard, the calculated concentration must deviate ≤ 10% from the nominal concentration. The analyte response at the lower limits of quantification and detection (LLOQ and LOD) must have a signal-to-noise ratio (S/N) ≥ 10 and three respectively. Plasma sample concentrations for the plasma half-life experiment after chronic exposure to 100 ppm SCN were run using the HPLC-UV method used in Breen et al. (2019) originally adopted from Rong et al. (2005). Samples were later confirmed using the UPLC-MS method detailed here.

### Statistics and analysis

The SCN half-lives and the accompanying regression statistics were determined using Origin 2018 (OriginLab, Northampton, MA, USA). The data were fit using the exponential fitting tool to a single exponential decay function, 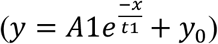, where x is time, y is concentration, y_0_ is the value of the function at the asymptotic limit, A1 is the concentration maximum, and t1 is the reciprocal of the first-order elimination rate constant k. None of the variables were constrained and no weighting function was used. Fits were to the full data set, the plasma concentration for each fish measured was treated as an individual sample point and were not averaged at each sampling interval prior to fitting. The reported half-life is related to the rate constant by 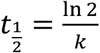.

## RESULTS

### Cyanide and Thiocyanate Exposure

Shortly after the fish were immersed in the 50 ppm CN bath, they began swimming erratically and gasping for air. Once in the recovery bath, a loss of balance and respiratory activity was observed. Both 20 s and 45 s exposed fish were completely immobilized by 45 seconds after initially being dipped into CN bath. Qualitatively recovery time varied proportionally to the exposure time. For the 20 s exposure, fish returned to normal swimming behavior by approximately 7 minutes while for the 45 s, normal swimming returned by approximately 17 minutes. There were no mortalities in any exposure. Most exposed fish did not accept food for the first 3 days post-exposure, but then ate regularly after the third day.

After exposure to CN, the SCN concentration in the plasma was observed to increase quickly over the first 6 hours, and then begin to decrease rapidly over the next 24 hours. The maximum SCN was observed 0.17 days post-exposure corresponding to a concentration of 468 ± 29 ppb for the 45 s exposure and 0.13 days post-exposure corresponding to a concentration of 301 ± 6 ppb for the 20 s exposure (Fig. 1A, B). During the course of our first CN exposure, the SCN in the plasma remained above the control level (SCN Conc. > 31 ppb ± 10) at our final data acquisition time of 13 days. In order to determine if and when SCN blood plasma levels reach control levels, a second exposure was carried out. In this second trial, the maximum SCN concentration of 399 ppb ± 60 was observed 0.55 days post-exposure, and control levels were reached seven days after exposure (Fig. 1C).

**Figure 1.**
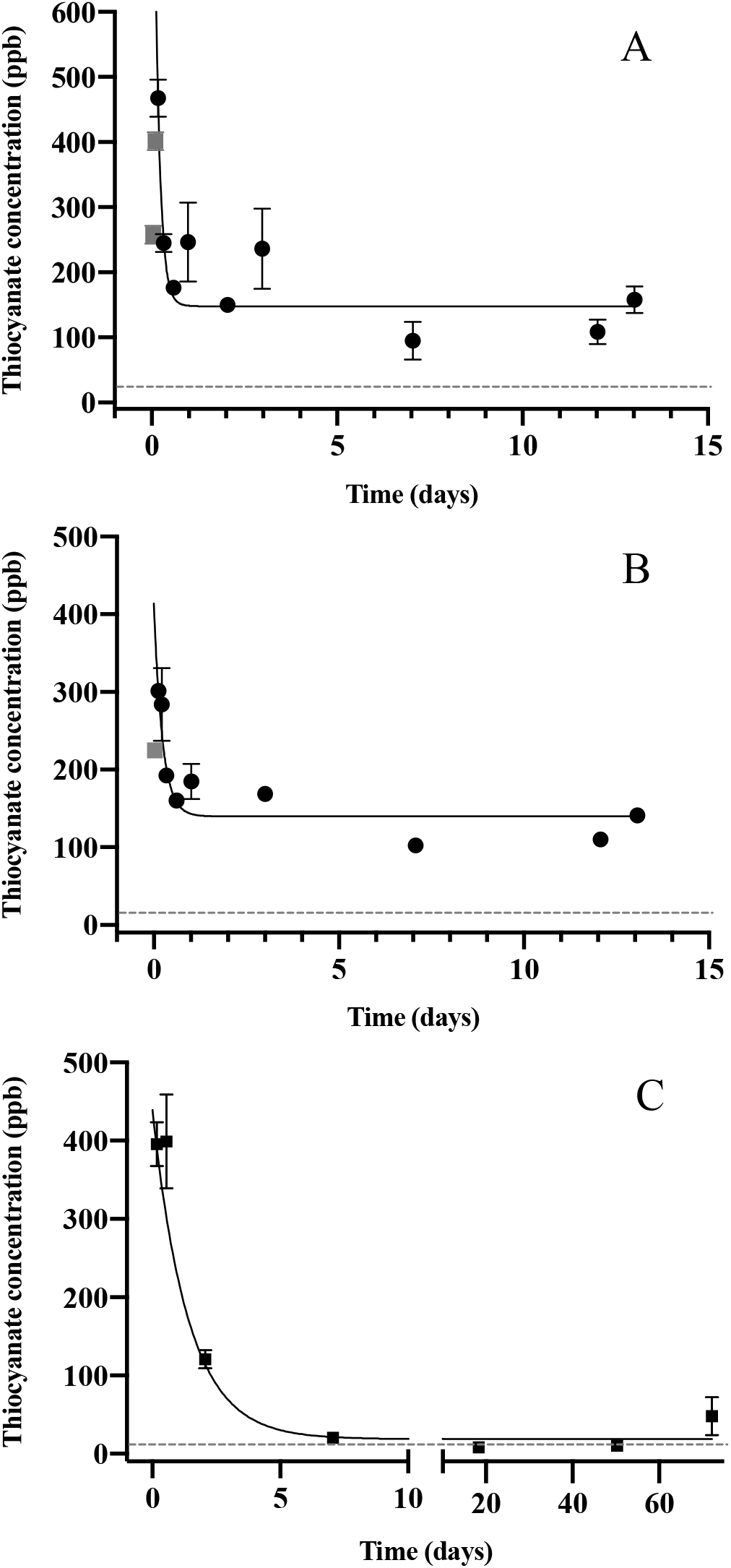
The SCN plasma concentration during depuration in *Amphiprion clarkii* after exposure to 50 ppm CN for trial 1 (A) 45 s and (B) 20 s with sampling out to 13 days and trial 2 (C) 45 s with sampling out to 72 days. The black solid line represents the fit of the data to a single-phase exponential decay function. Gray data points represent early time points where SCN plasma levels are increasing and are therefore not included in the fit. Each data point indicates the mean ± S.E. The dashed gray line represents the average SCN concentration (SCN Conc. > 34 ppb ± 10) in control fish. Where no error bar is observed the error is smaller than the data point.

In all three exposures, the rate of SCN elimination was fit to a single-phase exponential decay function with time constant parameters (Table 2). Regression statistics demonstrated all exposures to be statistically significant when fit to a single-phase exponential decay function (T1, 45 s: r^2^ = 0.578 and P < 0.0001, T1, 20 s: r^2^ = 0.676, P < 0.0001, T2: r^2^ = 0.86, P < 0.0001). The data were fit without constraining y_0_, and thus the concentration of SCN was not forced to go to zero at infinite time. This resulted in y_0_ values for the first trial of 163 ppb ± 15 for 45 s exposure and 136 ppb ± 10 for 20 s exposures which are above control levels and 7.8 ppb ± 17 for the second trial which is below control levels. In the first trial, the half-lives observed are fast, dropping to a plateau level in about two days following exposure. The plateau level remained for the next 13 days and no data were obtained for times longer than 13 days for this trial. In the second trial, the control levels were reached quickly, as no plateau was observed and the resultant half-life was determined to be 1.2 ± 0.2 days. In comparing the goodness of the fit from trial one to trial two, it appears that the trial two fit is much better than either of the trial one fits, based on the reported correlation coefficients and standard errors although there were fewer early time points sampled for the second trial.

**Table 2.** Half-life results of two acute CN exposure trials and one chronic SCN exposure with standard errors of the fit to the function 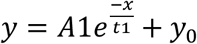 of the plasma SCN concentrations in *Amphiprion clarkii* exposed to 50 ppm CN or 100 ppm SCN. k and t_1/2_ are calculated from the fit parameter t_1_.

| Parameter | 45 s exposure,<br>trial 1 | 20 s exposure,<br>trial 1 | 45 s exposure,<br>trial 2 | SCN exposure |
| --- | --- | --- | --- | --- |
| $t_{1/2}$ (days) | $0.10 \pm 0.04$ | $0.20 \pm 0.06$ | $1.2 \pm 0.2$ | $0.13 \pm 0.02$ |
| $Y_0$ (ppb) | $163 \pm 15$ | $136 \pm 10$ | $7.8 \pm 17$ | $-580 \pm 1700$ |
| A1 (ppb) | $880 \pm 390$ | $261 \pm 54$ | $468 \pm 35$ | $55000 \pm 4000$ |
| $t_1$ (days) | $0.14 \pm 0.06$ | $0.29 \pm 0.09$ | $1.7 \pm 0.3$ | $0.19 \pm 0.03$ |
| k (days<sup>-1</sup>) | $6.9 \pm 2.7$ | $3.5 \pm 1.0$ | $0.58 \pm 0.09$ | $5.3 \pm 0.8$ |
| $r^2$ | 0.55 | 0.683 | 0.86 | 0.90 |
| P | P<0.0001 | P<0.0001 | P<0.0001 | P<0.0001 |
| N | 48 | 44 | 41 | 37 |

During the chronic SCN exposure (100 ppm, 12 days) experiments, fish behaved normally, ate well and showed no external signs of stress. Once depuration was initiated blood plasma levels were at a maximum (44 ppm ± 2.5) at the initial sample time (0.02 days) and decreased to average control levels (31 ppb ± 10) by day 15 (Fig. 2). Fitting of the data to a single-phase exponential decay resulted in a half-life of 0.13 ± 0.02 days with an r^2^ value of 0.90 (Table 2).

**Figure 2.**
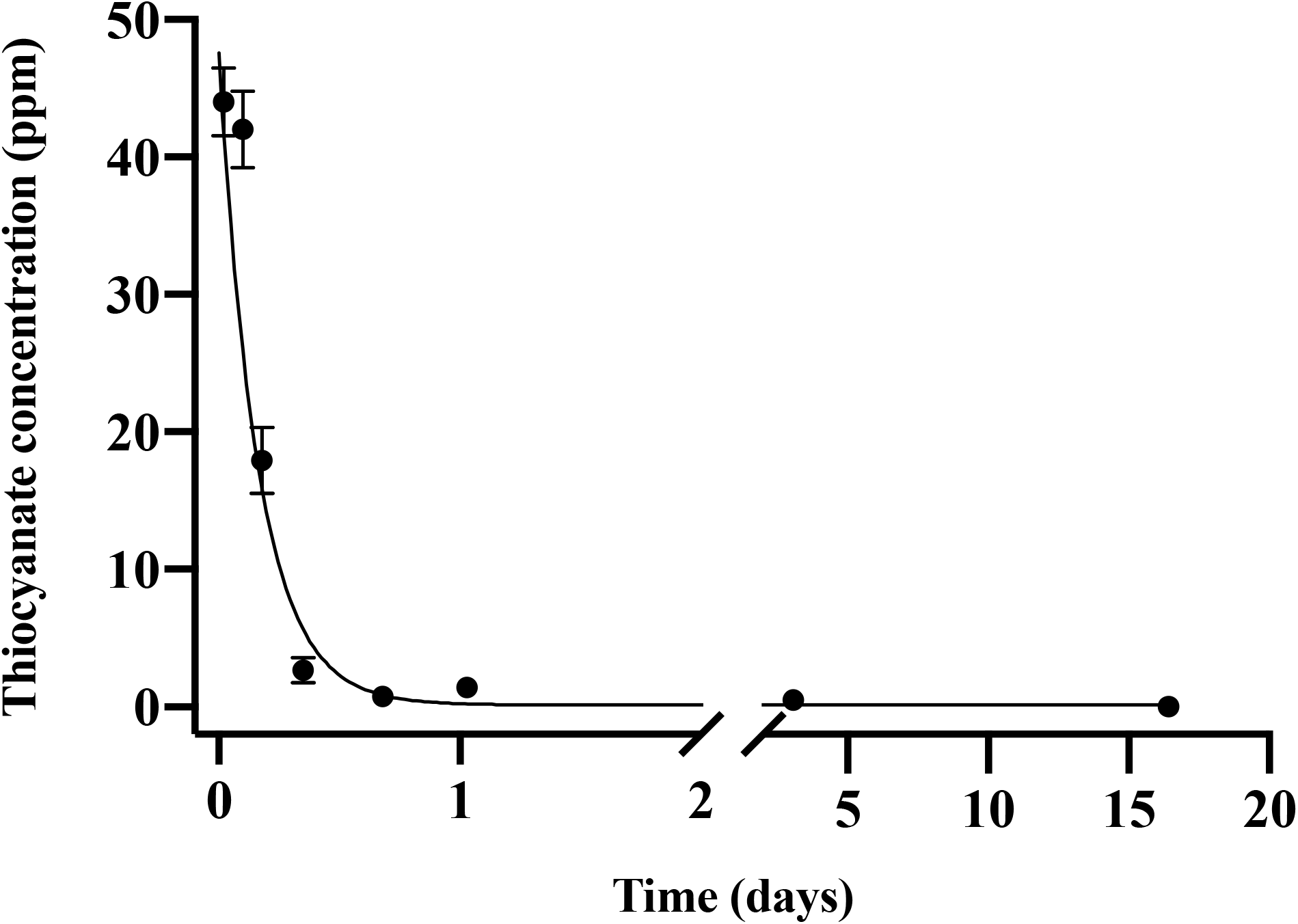
The plasma SCN concentration during depuration for *Amphiprion clarkii* after exposure to 100 ppm SCN for 12 days. The black solid line represents the fit of the data to a single-phase exponential decay function. Each data point indicates the mean ± S.E. Where no error bar is observed the error is smaller than the data point.

### Aquarium Water

After the first chronic exposure to 100 ppm SCN experiment, we were careful to rinse fish thoroughly before transferring to depuration tanks to remove SCN from the outer surface of the fish. In this chronic SCN exposure experiment, multiple fish were housed in 20 L buckets containing 15 L of saltwater, and this water was tested for the presence of SCN as a quick screen to see if any SCN was observable in the aquarium water. In this preliminary test, SCN levels were found to be in the 20-50 ppb range, at least for the first few hours, and SCN was only detected prior to the first water change. No SCN was detected in any of the aquarium water used as a check for contamination. After the 48 hour sampling the SCN blood plasma levels were 74 ppb ± 12. and then 26 ppb ± 12 after the last sampling at 72 hours. Because of this preliminary positive result, a second more systematic study of SCN levels in aquarium water during depuration from fish exposed to chronic levels of SCN was undertaken.

The results of the systematic aquarium water analysis for this chronic exposure to SCN are shown in Fig. 3. Water samples were collected at time zero just prior to and immediately after introducing a fish to its holding beaker. In these water samples, in both instances, no SCN was detected. At our next sampling point, the fish had been depurating for two hours and SCN levels were found to be at their highest level of 39 ppb ± 5. From this maximum, SCN concentrations decreased continuously and, when sampled at 24 hours before the first water change, no SCN was detected in the aquarium water. Sampling continued before daily water changes for three more days but no SCN in the aquarium water was detected at these later time points.

**Figure 3.**
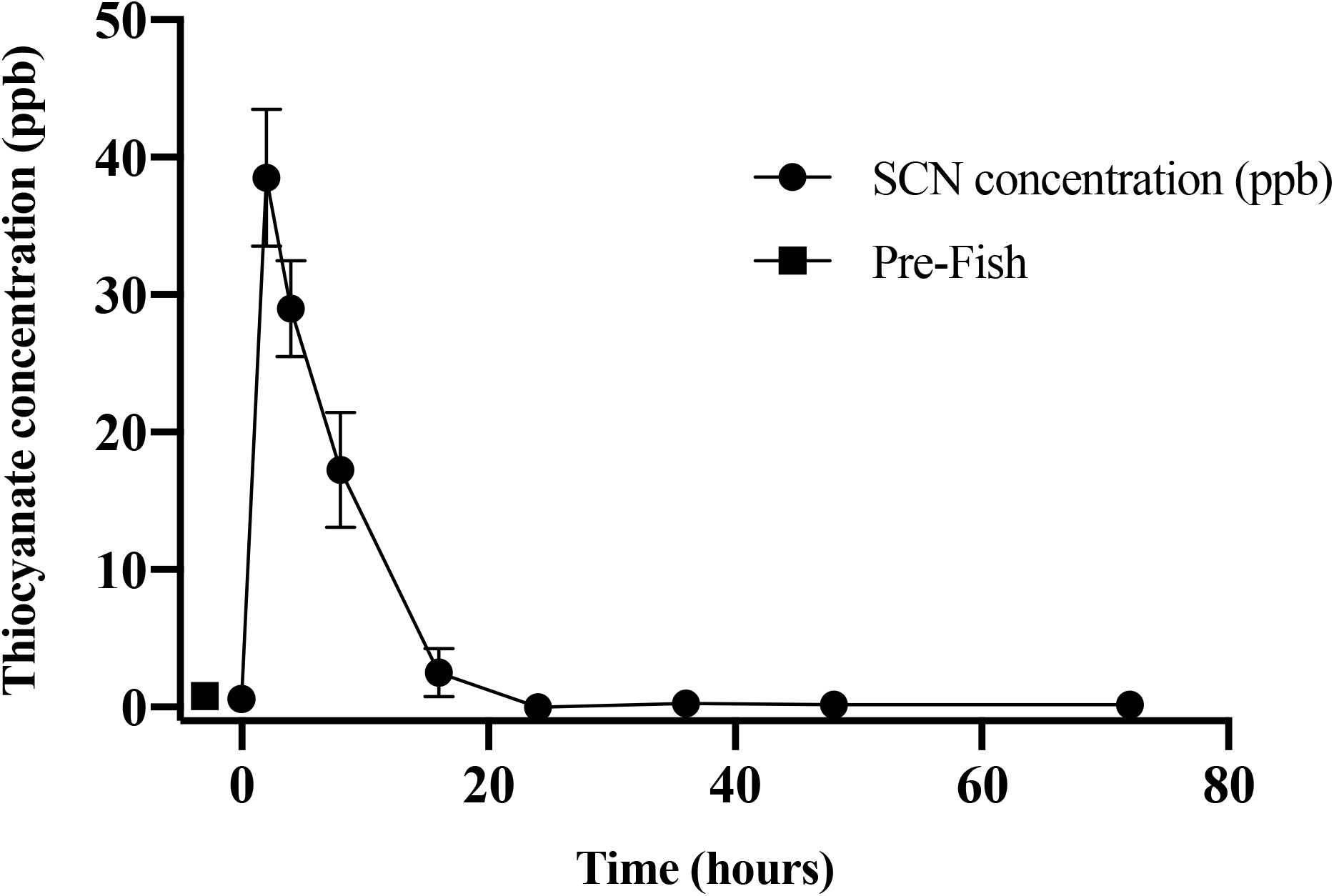
The average (n=10) concentration of SCN (ppb) of 15-L aquarium water, each containing a single *Amphiprion clarkii* depurating after exposure to 100 ppm SCN for 12 days. The data points indicate the mean ± S.E of the SCN concentration. Circles are samples once depuration had begun, the square indicates the measured SCN in the aquarium water prior to the addition of the fish. Where no error bar is observed the error is smaller than the data point.

This observation of an initial increase followed by an apparent loss of SCN in aquarium water over the first 24 hours of depuration after chronic SCN exposure required further study. Fish with no prior exposure to SCN were placed in beakers containing saltwater spiked with SCN and sampled for 24 hours. The concentration of SCN measured in the aquarium water with a single *A. clarkii* held in 500 mL of saltwater continually decreased from 15 ppb initially to below our detection limit at 16 hours (Fig. 4). After the first water change at 24 hours, the fresh saltwater was again spiked with SCN and the SCN concentration was found to be 15 ppb ± 1. However, once again when sampled at 24 hours the SCN concentration had dropped to below our detection limit (1 ppb). After the 24 hour sampling the SCN blood plasma levels were 429 ppb ± 164 and remained 429 ppb ± 94 after the last sampling at 48 hours.

**Figure 4.**
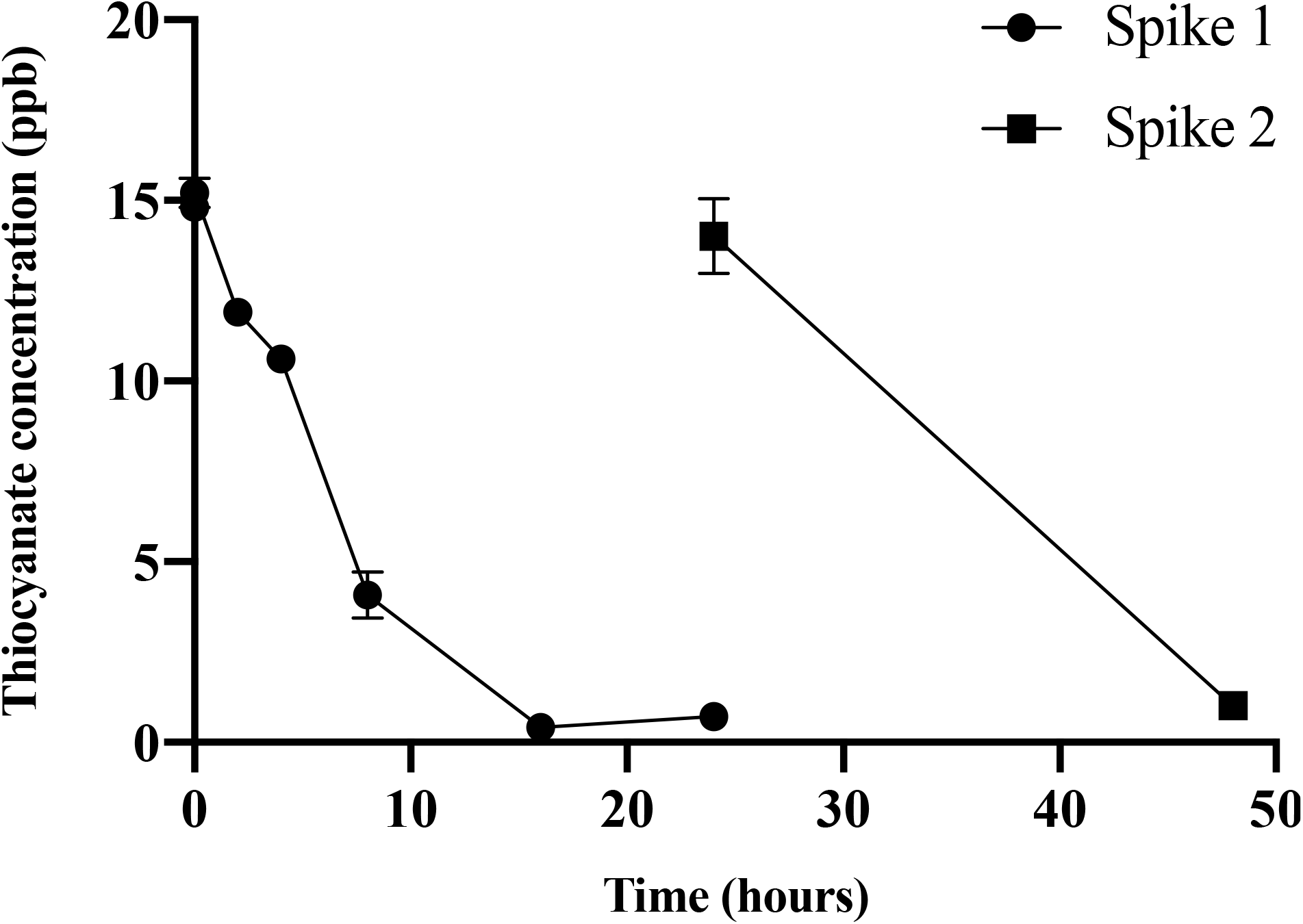
Average (n=10) SCN concentration of 500 mL aquarium water spiked with SCN containing 1 non-exposed *Amphiprion clarkii*. Each data point indicates the mean ± S.E. Where no error bar is observed the error is smaller than the data point. The round data points indicate the data collected after the first SCN spike and the square data points show that data after a water change and a second 20 spike.

## DISCUSSION

### CN Exposure

*Amphiprion clarkii* exhibited similar behavior as *A. ocellaris* when exposed to CN. Erratic behavior was followed by a loss of equilibrium and paralysis. In comparing these results with our previously reported work on *A. ocellaris,* the maximum level of SCN in blood plasma was decidedly lower for *A. clarkii* than for *A. ocellaris* while the half-lives reported were similar or smaller for *A. clarkii* depending on the exposure time (Table 3). The lower concentration of SCN in blood plasma and faster half-life suggest that *A. clarkii* take up less CN during exposure when compared to *A. ocellaris*. The half-lives reported here for marine fish are in reasonable agreement with those reported for mammals (0.21-8.3 days) (Logue et al. 2010).

**Table 3.** Summary of SCN half-lives in fish exposed to CN or SCN including relative standard deviation (RSD) and average fish weight. Summary of max concentration of SCN observed in fish exposed to CN or SCN.

| Model species |  | CN 20 s<br>trial 1 | CN 45 s<br>trial 1 | CN 45 s<br>trial 2 | SCN<br>chronic | Reference |
| --- | --- | --- | --- | --- | --- | --- |
| *Amphiprion*<br>*ocellaris* | Half-life<br>(days) | 0.44 ± 0.15 | 1.01 ± 0.26 | NA | 0.35 ± 0.07 | (Breen et al.<br>2019) |
|  | RSD | 34% | 26% |  | 20% |  |
|  | Mass<br>(g) | 5.68 ± 1.42 | 5.03 ± 1.41 |  | 3.84 ± 0.59 |  |
|  | Max. Conc<br>(ppm) | 1.9 ± 0.6 | 2.3 ± 0.2 |  | 220 ± 31 |  |
| *Amphiprion clarkii* | Half-life<br>(days) | 0.20 ± 0.06 | 0.10 ± 0.04 | 1.2 ± 0.2 | 0.13 ± 0.02 | Current study |
|  | RSD | (30%) | (40%) | (17%) | (15%) |  |
|  | Mass<br>(g) | 3.07 ± 0.62 | 3.54 ± 0.78 | 12.84 ±<br>9.37 | 6.09± 2.72 |  |
|  | Max. Conc<br>(ppm) | 0.301±<br>0.066 | 0.468 ±<br>0.028 | 0.399 ±<br>0.060 | 44 ± 2.5 |  |
| *Oncorhynchus*<br>*mykiss* | Half-life<br>(days) | NA |  |  | 2.02 ± 0.06 | (Brown et al.<br>1995) |
|  | RSD |  |  |  | 3% |  |
|  | Mass<br>(g) |  |  |  | 20 |  |
|  | Max. Conc<br>(ppm) |  |  |  | 60.5 ± 6.2 |  |

For our first CN exposure trial the half-life of the 45 s exposure was faster than the 20 s exposure, contrary to those results for *A. ocellaris.* However, both of these half-lives have large RSDs (40%, 30%), and the argument could be made that there is no difference within their respective uncertainties. Our second 45 s trial, which has a much lower RSD than the first 45 s trial (17% vs. 40%) has a half-life similar to that observed for *A. ocellaris*. The goal of the second trial was to establish when control levels were reached, and because of this, there were fewer early time points sampled. Trial 1 CN exposures have more sampling data points at early time points, resulting in a better fit of the elimination from the blood plasma rather than the terminal half-life.

SCN concentration for *A. ocellaris* plateaued between two and four days at approximately 500 ppb for both 20 and 45 s exposure times (Breen et al. 2019). In the first trial of the current study, the SCN concentration plateaued between four and 12 hours at approximately 150 ppb for both exposure times, likely due to the faster half-life and likely lower dose of CN ingested.

Vaz et al. (2012) exposed *A. clarkii* (1.8 ± 0.2 g) to 25 ppm CN for 60 seconds and reported a 33% mortality rate. We did not observe any mortality when fish were exposed to CN concentrations of 50 ppm CN for 45 seconds. However, this difference in vulnerability could be due to differences in fish size, health or stress level (Hanawa et al. 1998). When assessing the vulnerability of *A. ocellaris* to CN, Madeira et al. (2020) found that larger fish had a higher survival rate and quicker recovery time when exposed for the same time and concentration of CN as their smaller conspecifics. As the fish used in this study were larger, a lower mortality rate would be expected.

### SCN Exposure

As with the CN exposure, when the results for *A. clarkii* are compared with those previously reported for *A. ocellaris*, the SCN blood plasma level was much lower. Both species were exposed to 100 ppm SCN bath for 11-12 days, but the maximum SCN blood plasma was 44 ppm ± 2.5, while that for *A. ocellaris* was at 220 ppm ± 31 (Table 3). The SCN levels in the blood plasma of *A. clarkii* were half that of the exposure bath, while those observed for *A. ocellaris* were twice that of the exposure bath. It appears that even under different exposure conditions (pulsed CN versus chronic SCN), *A. clarkii* have lower levels of SCN in their bloodstream, indicating that they are up-taking less CN/SCN during exposure than *A. ocellaris* or eliminating SCN from their blood more efficiently.

The half-life for SCN clearance in the blood plasma measured for *A. clarkii* when chronically exposed to SCN is 63% faster when compared to *A. ocellaris*. In a similar experiment on the freshwater fish rainbow trout *(Oncorhynchus mykiss*) the reported half-life for clearance of SCN from their blood was 2.02 or 2.36 days depending on the model used (Brown et al. 1995). This is much slower than our reported value of 0.13 ± 0.02 days and could reflect the differences in the osmoregulatory systems of marine versus freshwater fish.

### Aquarium Water

Failure to replicate the work of Vaz et al. (2012) calls to question the ultimate fate of CN and SCN in marine fish. We have now confirmed in two marine species that CN is converted to SCN quickly as evidenced by the rapid rise of SCN in the blood plasma following CN exposure in accordance with the mammalian model. The clearance rate of SCN from the blood plasma is also quick, with the highest levels depleted in a few days after CN exposure, also in accordance with mammalian models. However, the question remains where does the SCN go? Is the failure to detect SCN in the aquarium water of marine fish post-acute exposure to CN because they are not excreting it, or is it because any excretion by a small fish in a relatively large quantity of water is diluted to concentrations below the detection limit?

Blood plasma levels of SCN following acute exposure to CN were found to be in the range 0.4 – 2 ppm at their maximum and only for a day or two after exposure. Levels above this are unlikely as higher CN doses would be required, leading to an increase in mortality rather than higher levels of SCN in the blood plasma. In order to enhance detection capabilities by increasing the SCN in the blood plasma, which presumably would lead to higher levels of SCN in the urine if dilution was the limiting factor, *A. clarkii* were exposed to 100 ppm SCN for 12 days, presumably completely saturating the fish. Under such conditions, blood plasma levels were found to be close to 50 ppm for *A. clarkii*, approximately 25 times higher than the highest SCN concentration observed in CN exposed fish. These fish were placed in 500 mL of aquarium water to depurate, a volume three times smaller than the 1.5 L used by Vaz et al. (2012). It was believed that higher SCN blood plasma levels as compared to CN exposure and the smaller aquarium water volume used in depuration should increase the likelihood of observing SCN in the aquarium water. In fact, SCN was detected in the aquarium water of the depurating fish, a maximum was observed close to 40 ppb two hours after depuration began. However, after this initial observation, SCN concentration steadily decreased until the first water change at 24 hours where it reached LOD. Sampling continued before each water change for three days, but no further SCN was detected in the aquarium water. These results demonstrate that *A. clarkii* have the capability of absorbing low levels of SCN, likely through their drinking response. We speculate that the source of the SCN we detected two hours after introducing the fish to the beaker is possibly due to diffusion out of the slime coat and not via urinary excretion, as no other SCN was observed at later times. Once the SCN is introduced to the aquarium water from the slime coat, it is then absorbed by the fish via the drinking response. However, drinking rates in marine fish have been reported to range from 2-7 mL/kg per hour making it unlikely for the fish to drink 500 mL of aquarium water in less than 24 hours. (Perrott et al. 1992; Fuentes & Eddy 1997; Grosell 2019).

Given that the time scale (~ 20 hours) for SCN blood plasma clearance and for SCN aquarium water clearance are similar (Fig. 2, Fig. 3), it is also possible that most of the SCN absorbed in the chronic exposure may have been rapidly excreted. Then after the initial 24 hours, further excretion is too low to detect or the rate of reabsorption exceeds that of excretion for the following three days. However, this scenario is less likely since SCN is continually and rapidly eliminated from the blood plasma for the first 12 hours of depuration, not just the first two hours.

Our results testing SCN in aquarium water is the reverse of those of Vaz et al. (2012). In their study, they did not observe any SCN in aquarium water in the first 24 hours post-acute exposure to CN but then observed a steady increase of SCN concentration in aquarium water up until 28 days post-exposure. Our results show that there was no continual SCN excretion even after chronic exposure to SCN, but rather, low-level absorption was observed. This observation was unexpected but was further confirmed in the subsequent experiment where fish were placed in aquarium water that was spiked with SCN.

It has been previously speculated that SCN was excreted out of the fish via urine following CN exposure (Vaz et al. 2012) into the surrounding water to eliminate the ion from the body as in mammalian models (Lanno & Dixon 1996; Nelson 2006; Logue et al. 2010). However, the data presented here suggest that, not only is SCN not excreted by *A. clarkii*, but that the fish will uptake low doses of SCN from the holding water and retain SCN. This further corroborates the findings that testing for SCN in the holding water of this fish is not a viable indicator of CN exposure (Herz et al. 2016; Breen et al. 2018), opposing the previous studies (Vaz et al. 2012; Vaz et al. 2017). This also nullifies any concern of false positives from non-exposed fish up-taking SCN during cohabitation with exposed fish excreting SCN in aquarium water. Howerver, this may raise concern of fish uptaking SCN in aquarium water contaminated with SCN from other sources.

This difference in SCN metabolism compared to mammalian models may be due to the osmoregulation strategy of marine fish. To maintain osmotic balance with the surrounding water, marine fish have a relatively low urinary excretion rate in order to retain water in their blood. Urine flow rates in marine teleosts are minimal at 1–2% of body weight daily because water is highly conserved and can be reabsorbed by the urinary bladder (Evans 1993). This lends further support to the claim that any SCN in the aquarium water during depuration after chronic exposure to SCN was likely due to its release from the slime coat of the fish. Most of the components in the saltwater that fish ingest are absorbed by the esophagus and intestine. Certain excess ions such as Na^+^, Cl^−^, and K^+^ that are absorbed by the gut are expelled from the body by specialized, mitochondria-rich cells called chloride cells (Greenwell et al. 2003). The divalent ions such as Mg^2+^ and SO_4_^2−^ are excreted in the feces (Hickman Jr 1968). However, the excretion pathway or terminal fate of SCN ions in the body of marine fish remains unknown.

The results of the *A. clarkii* half-life assessments indicate a two-compartment model for SCN elimination similar to that of *A. ocellaris* (Toutain & Bousquet-mélou 2004). Two-compartment models are commonly used in pharmaceutical research for drug metabolism (Metzler 1971). As CN is absorbed through the gills and/or the gut via the drinking response, it is metabolized into SCN, and the SCN concentration increases in the blood plasma of the fish (Breen et al. 2019). We observed that SCN is rapidly eliminated from the plasma. SCN blood plasma levels dropped while depurating for 72 hours after exposure to 100 ppm SCN and stayed the same after being introduced to SCN spiked aquarium water after 24 hours and 48 hours after a second spike after the 24 hour sampling. Aquarium water data demonstrates that SCN is not leaving the fish, therefore it may be entering another compartment within the fish’s body. The fish’s tissue may be acting as a SCN reservoir.

SCN stored in the tissue of fish could also serve as a potentially longer post-exposure indicator of CN exposure but further research on the half-life of SCN in the tissue of marine fish exposed to CN would need to be conducted. This study provides a good starting point for discussion and further research.

In order to properly validate a CN detection method utilizing SCN, it would also be critical to know the baseline values of SCN in the blood and tissue of non-cyanide caught fish from the areas where CN is likely used. Fish in these areas may be exposed to very low concentrations of CN from natural sources such as cyanogenic foods therefore already containing SCN in blood and tissues. Knowing this information will reduce the potential for false positives. The present findings confirm the blood plasma SCN maybe be a useful biomarker of CN exposure in marine fish if measured shortly after exposure. Additional species from a broader taxonomic sample must be evaluated prior to any definitive conclusions.

## ACKNOWLEDGEMENTS

This work was funded in part by the Pet Industry Joint Advisory Council. We would like to thank the current and former undergraduates from Roger Williams University who have been involved in this project especially, Julia Grossman, Gabbie Baillargeon, Hannah Sterling, Natalie Danek, Julia Dwyer, and Sara Hunt. Kevin Erickson of MASNA provided thoughtful comments on an earlier draft of this manuscript. Sea & Reef Aquaculture of Maine U.S.A. provided a portion of the fish used in these experiments.

